# Dynamic cholinergic tone in the basal forebrain reflects reward-seeking and reinforcement during olfactory behavior

**DOI:** 10.1101/2020.11.30.404798

**Authors:** Elizabeth Hanson, Katie L Brandel-Ankrapp, Benjamin R Arenkiel

**Affiliations:** Department of Molecular and Human Genetics, Baylor College of Medicine, Houston, TX, USA; Postbaccalaureate Research Education Program, Baylor College of Medicine, Houston, TX, USA

**Keywords:** acetylcholine, olfaction, basal forebrain, go/no-go, reward, discrimination, GABA, top-down

## Abstract

Sensory perception underlies how we internalize and interact with the external world. In order to adapt to changing circumstances and interpret signals in a variety of contexts, sensation needs to be reliable, but perception of sensory input needs to be flexible. An important mediator of this flexibility is top-down regulation from the cholinergic basal forebrain. Basal forebrain projection neurons serve as pacemakers and gatekeepers for downstream neural networks, modulating circuit activity across diverse neuronal populations. This top-down control is necessary for sensory cue detection, learning, and memory, and is disproportionately disrupted in neurodegenerative diseases associated with cognitive decline. Intriguingly, cholinergic signaling acts locally within the basal forebrain to sculpt the activity of basal forebrain output neurons. To determine how local cholinergic signaling impacts basal forebrain output pathways that participate in top-down regulation, we sought to define the dynamics of cholinergic signaling within the basal forebrain during motivated behavior and learning. Towards this, we utilized fiber photometry and the genetically encoded acetylcholine indicator GAChR2.0 to define temporal patterns of cholinergic signaling in the basal forebrain during olfactory-guided, motivated behaviors and learning. We show that cholinergic signaling reliably increased during reward-seeking behaviors but was strongly suppressed by reward delivery in a go/no-go, olfactory-cued discrimination task. The observed transient reduction in cholinergic tone was mirrored by a suppression in basal forebrain GABAergic neuronal activity. Together, these findings suggest that cholinergic tone in the basal forebrain changes rapidly to reflect reward-seeking behavior and positive reinforcement to impact basal forebrain circuit activity.

## 1 Introduction

Rapid and precise sensory processing is critical for properly interpreting the external world. As a chemical sense, olfaction requires the ability to sample a vast, non-continuous sensory space with a wide range of stimulus intensities (Ache and Young, 2005). For this, the olfactory system must quickly separate and identify trace amounts of volatilized signals from a complex, noisy background (Rokni et al., 2014). However, as an animal moves through the world, the contexts in which it encounters odors, as well as its own internal drives, are constantly changing. Therefore, olfactory processing must be flexible as well as sensitive in order to facilitate these changing needs. Flexible olfactory processing depends, in part, on top-down regulation (Restrepo et al., 2009; Pashkovski et al., 2020). Top-down regulation is a feature of sensory systems through which information about an animal’s context, internal state, or previous experience modulates circuit function to sculpt the way stimuli are perceived (Gilbert and Sigman, 2007). In olfaction, for example, top-down regulatory mechanisms are recruited during active sensing in ways that improve odor detection and discrimination (Jordan et al., 2018), allow odor detection within a single sniff (Laing, 1986; Rinberg et al., 2006), and facilitate adaptive filtering during high frequency bouts of sniffing (Verhagen et al., 2007). Top-down regulation also allows for rapid changes in odor responses depending on context (Kay and Laurent, 1999; Beshel et al., 2007; Kudryavitskaya et al., 2020), and directly influences plasticity within the olfactory system (Fletcher and Wilson, 2003; Fletcher and Chen, 2010; Lepousez et al., 2014; Hanson et al., 2020).

An important source of top-down regulation in olfaction comes from the horizontal limb of the diagonal band of Broca (HDB) in the basal forebrain (Zaborszky et al., 1986; Mandairon et al., 2006; Gracia-Llanes et al., 2010; Ma and Luo, 2012; Rothermel et al., 2014). Basal forebrain neurons mediate state-dependent top-down regulation through signaling mechanisms that span diverse time scales ranging from milliseconds to hours (Buzsaki et al., 1988; Détári et al., 1999; Muñoz and Rudy, 2014). Fast, phasic signals from the basal forebrain mediate effects of attention on sensory processing, decision making, and sensory cued task performance (Parikh et al., 2007; Lin and Nicolelis, 2008; Pinto et al., 2013; Muñoz and Rudy, 2014; Hangya et al., 2015; Gritton et al., 2016). It has long been hypothesized that basal forebrain cholinergic signaling in particular mediates attentional effects on sensory processing circuits (Mandairon et al., 2006; Herrero et al., 2008; Chaudhury et al., 2009; Ghatpande and Gelperin, 2009; Goard and Dan, 2009; Ma and Luo, 2012; Chapuis and Wilson, 2013; Zhan et al., 2013; Rothermel et al., 2014). However, it has also been found that that non-cholinergic neuronal activity better predicts behavioral variables associated with attention in an auditory-cued go/no-go task (Hangya et al., 2015). Additionally, a recent study has described anticipatory activity among both cholinergic and non-cholinergic neurons in the basal forebrain during an olfactory-cued go/no-go task (Nunez-Parra et al., 2020). Importantly, in agreement with these earlier studies (Dannenberg et al., 2015; Xu et al., 2015), cholinergic neurons were also noted to collateralize within the basal forebrain to influence the activity of neighboring non-cholinergic neurons during task performance (Nunez-Parra et al., 2020). Together, this evidence suggests that non-cholinergic basal forebrain neurons mediate effects of attention on sensory processing, and it raises the question of how communication between cell types within the basal forebrain controls statedependent basal forebrain output.

Parallel cholinergic and GABAergic projections from the basal forebrain to the olfactory bulb mediate distinct features of top-down regulation (Böhm et al., 2020). Separately, the cholinergic and GABAergic projections control gain, signal-to-noise ratio, habituation, and oscillatory activity, and odor discrimination (Ma and Luo, 2012; Nunez-Parra et al., 2013; Rothermel et al., 2014; Ogg et al., 2018; Villar et al., 2020). Though both types of basal forebrain projections are important modulators of olfactory bulb odor and sniff responses, the upstream mechanisms that control basal forebrain output remain largely unknown. Ultimately, understanding how the basal forebrain mediates state-dependent changes in olfactory processing requires a more detailed knowledge of signaling within the HDB, and how it controls HDB output during olfaction and complex olfactory-guided behavior.

Here we describe temporal patterns of cholinergic signaling within the HDB during an olfactory-cued go/no-go discrimination task where mice learn to associate one of two odors with a reward. Historically, monitoring acetylcholine directly, *in vivo*, with high temporal resolution, has been challenging. However, with the advent of the genetically encoded GPCR Activation-Based (GRAB) fluorescent sensor for acetylcholine (GACh2.0), we directly recorded rapid fluctuations in acetylcholine levels from freely moving, behaving animals (Jing et al., 2018). Combining targeted sensor expression with implanted fiber optics and fiber photometry, we directly recorded acetylcholine signaling from the basal forebrain chronically, during freely moving behavior. We found that acetylcholine levels within the basal forebrain are dynamic and bidirectionally regulated during performance of the go/no-go discrimination task. Reward-seeking behavior reliably evoked rapid increases in HDB acetylcholine, while positive feedback transiently suppressed cholinergic tone. These dynamics suggest that local cholinergic signaling rapidly modulates HDB circuitry, potentially mediating moment-to-moment changes in projection output and HDB-mediated top-down regulation in olfaction.

## 2 Results

### 2.1 Fiber photometry of a genetically encoded acetylcholine sensor reveals real-time cholinergic signaling in the basal forebrain

Defining the temporal profile of basal forebrain cholinergic signaling during complex behavior is a necessary step in determining how local cholinergic signaling impacts HDB circuit function and state-dependent output. To directly monitor cholinergic signals within the basal forebrain we injected wildtype mice with an adeno-associated virus (AAV) engineered to drive pan-neuronal expression of the acetylcholine sensor GACh2.0 (AAV hsyn-GACh). At the same time, we implanted a fiberoptic over the HDB (Figure 1A). Expression and targeting were verified *post hoc* in all mice via immunofluorescence and histology (Figure 1B). After implantation and injection, mice were allowed to recover and given three weeks to express the sensor prior to photometric recordings (Figure 1C). We first examined cholinergic signaling in freely moving mice during exploration of an open field arena. For this, we video-recorded mice exploring an open field while simultaneously using fiber photometry to record activitydependent changes in GACh2.0 fluorescence in the HDB (Figure 1C, D). During open field exploration, we observed both excitation and suppression events (Figure 1E). Notably, the detected events were not correlated with motion or position in the open field (Figure 1F), and the amplitude of the fluorescence signals (dF/F) were not correlated with speed (cm/s) over time (Pearson’s correlation = −0.043 ± 0.025, N = 4 sessions, 4 animals). These results revealed frequent spontaneous cholinergic signaling events in the HDB during behavior, which was not triggered by, or directly correlated with, voluntary locomotion.

**Figure 1:**
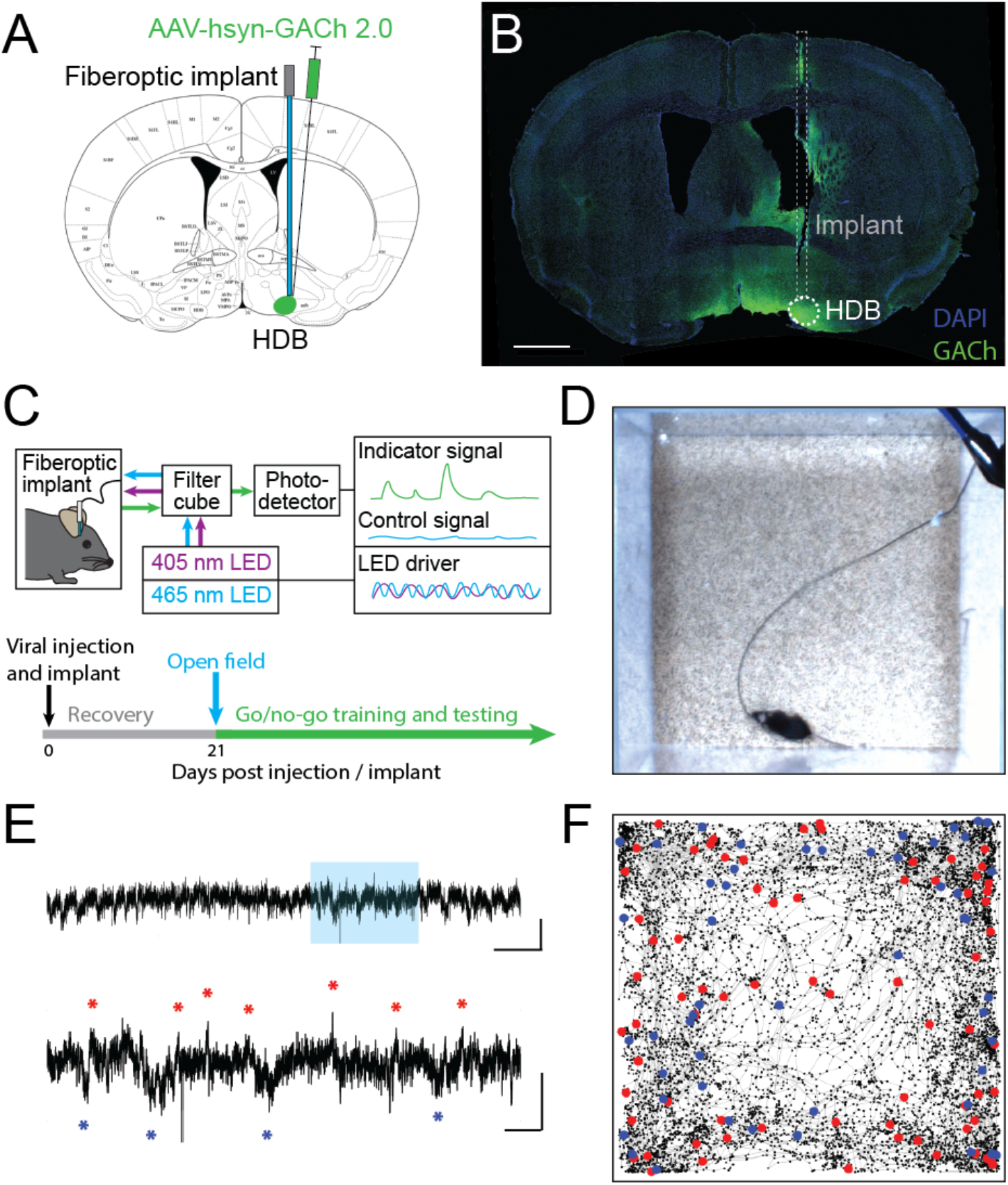
Fiber photometry of a genetically encoded acetylcholine sensor in the HDB reveals real-time cholinergic signaling during freely moving behavior. **A**. Coronal section schematic showing AAV injection and implant targeting the HDB. **B.** IHC of a coronal section showing GACh2.0 expression and implant targeting in the HDB. Scale bar = 1 mm. **C.** Schematic of photometry system showing light paths, LED control systems, filtering, and photodetection. Timeline of surgery, recovery, and photometry recording during open field exploration, followed by behavioral training and testing. **D.** Still frame from video of open field exploration with photometric recording. **E.** (Top panel) Isosbestic-subtracted GACh dF/F trace during open-field arena exploration. Y scale bar = 1 dF/F and X scale bar = 60 s. (Bottom panel) zoom of blue shaded portion of trace in top panel with excitation events marked with red asterisks and suppression events marked with blue asterisks. Y Scale bar = 1 dF/F, X scale bar = 10 s. **F.** Track of mouse location over 20 minutes of open field exploration with locations corresponding to increases in HDB cholinergic signaling (excitation events) marked with red dots and decreases (suppression events) marked with blue dots.

### 2.2 Basal forebrain cholinergic signaling rapidly fluctuates with reward-seeking and positive reinforcement

If HDB cholinergic signaling influences state-dependent basal forebrain output, we reasoned that the cholinergic reporter responses may dynamically change with behavioral states during the performance of complex, olfactory-guided, operant behaviors. To test this, we recorded photometry signals from freely moving mice performing an olfactory-cued go/no-go discrimination task (N = 33 sessions, 6 animals) (Figure 2A, B). Mice first underwent a shaping period of 10-14 days where they learned the mechanics of the task without photometry recording. During shaping, mice were trained to self-initiate trials by poking their nose into a port where they were presented with one of two odors. They then learned to distinguish between the delivery of an S+ odor, which indicated the availability of a water droplet at a separate reward port, and an S− odor, which indicated that no reward was available. A correct response to the S+ odor where a reward was obtained was considered a “Hit”. A correct response to the S− odor where a new trial was initiated without reward-seeking was considered a “Correct Reject”. An incorrect attempt to seek a reward after the S− odor was considered a “False Alarm” and an incorrect trial re-initiation after presentation of the S+ odor was considered a “Miss” (Figure 2A). Notably, this freely moving go/no-go task did not include punishment in response to False Alarms. Thus, feedback during odor-association learning was limited to positive reinforcement of a water reward in Hit trials, and negative reinforcement of a 4-second timeout after false alarms. Another feature of the freely moving task was that animals were required to self-initiate trials and reward-seeking. Thus, the timing of trial initiation and reward-seeking was determined entirely by the mouse, and it required both active engagement with the task and locomotion (Figure 2B).

**Figure 2:**
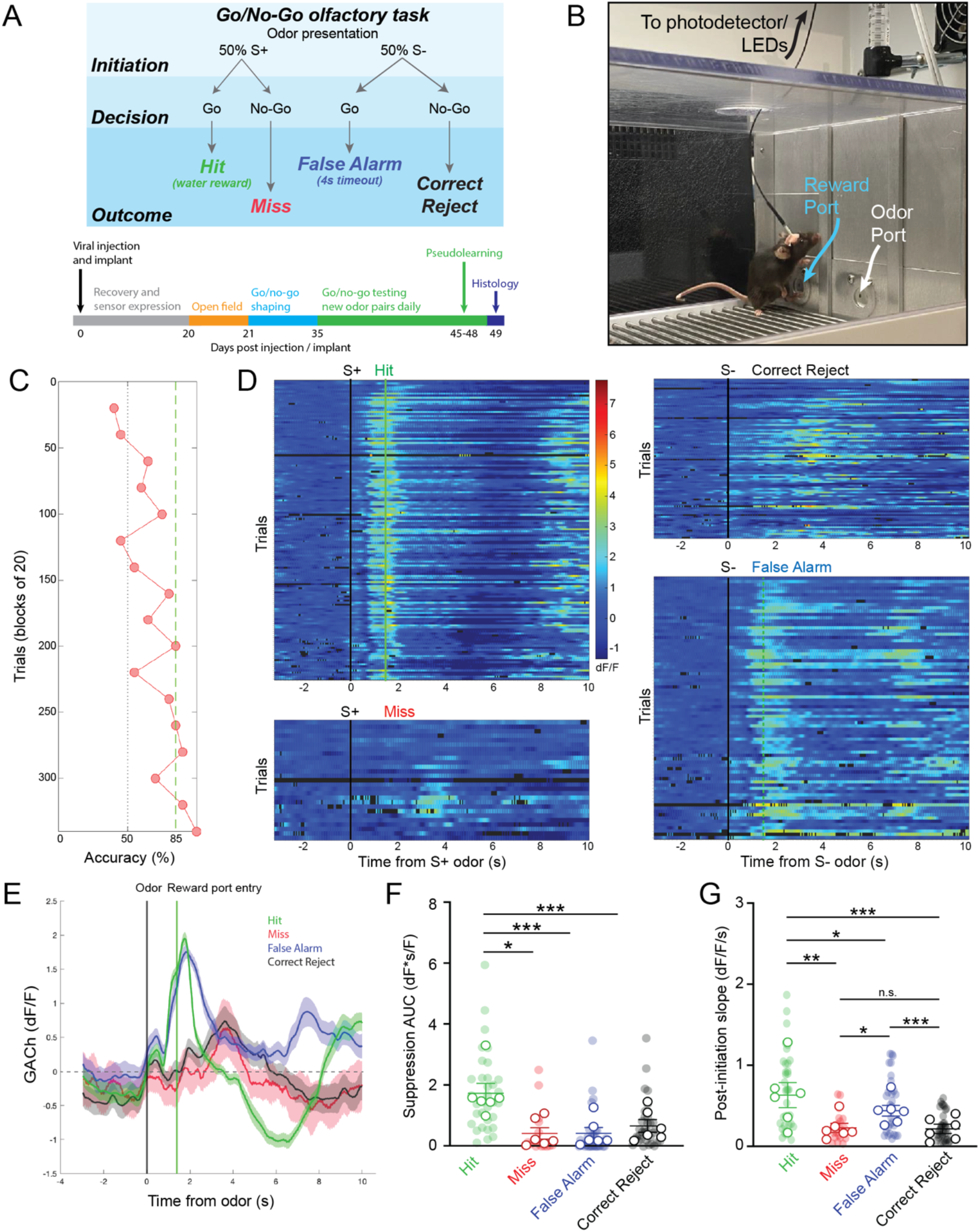
Basal forebrain cholinergic signaling reflects reward-seeking and positive reinforcement in an olfactory-cued go/no-go discrimination task. **A.** (Top panel) Schematic of olfactory-cued go/no-go discrimination task showing odor presentation, decisions, and possible trial outcomes (Hit, Miss, False Alarm, and Correct Reject). (Bottom panel) Timeline of surgery, recovery, and behavioral shaping and testing with photometry recording. **B.** Picture of mouse performing go/no-go task during photometric recording. **C.** Accuracy in blocks of 20 trials for a go/no-go testing session with novel odors highlighting chance (50%) and criteria (85%) levels. Accuracy is from the same session as the trials shown in D and E. **D.** Heatmap showing isosbestic-subtracted GACh dF/F from individual trials in a single go/no-go testing session. Trials are aligned by trial initiation time and divided by trial outcome. **E.** Average GACh dF/F traces for each trial type in the session shown in D. Shaded areas represent 95% confidence intervals. Black line marks trial initiation time. Green line marks the average reward port entry time in Hit and False Alarm trials. **F.** Area under the curve of suppression below baseline across trial types and testing sessions. Transparent circles represent individual testing sessions. Hollow circles represent mean values from all sessions completed by individual mice. Lines and error bars show mean ± SEM of means from each animal. *p < 0.05, *** p < 0.001 two-way nested repeated measures ANOVA with Tukey correction for multiple comparisons. **G.** Slopes of GACh dF/F after trial initiation across trial types and testing sessions. Transparent circles represent individual testing sessions. Hollow circles represent mean values from all sessions completed by individual mice. Lines and error bars show mean ± SEM of means from each animal. *p < 0.05, **p < 0.01, *** p < 0.001 two-way nested repeated measures ANOVA with Tukey correction for multiple comparisons.

Following the shaping period, we next recorded photometric signals from the basal forebrain while mice learned to discriminate novel odor pairs. As mice learned new odor pairs, success rates (accuracy within a block of 20 trials) increased. An odor pair was considered “learned” after two consecutive trial blocks with greater than 85% success (Figure 2C). Once proficient at the task, mice typically learned new odor-reward associations within a single training session of 200-300 trials (Trials to learn = 104.6 ± 8.3, N = 56 sessions, 12 animals). Using the timing of IR beam breaks at the odor port and reward port, individual trials of the go/no-go task were segmented into periods before and after trial initiation and, in the case of Hit and False Alarm trials, before and after reward-seeking. Aligning trials by initiation times, and separating them by trial type, revealed distinct temporal patterns of cholinergic signaling in each trial (Figure 2D). Averaging across trials showed that bidirectional changes in HDB cholinergic signaling were consistent within trial types, but distinct across trials (Figure 2E).

A notable feature of the signal specific to Hit trials was the suppression of HDB cholinergic tone following reward delivery. The magnitude of the suppression, calculated as the area under the curve (AUC), was significantly larger in Hit trials (1.73 ± 0.33 dF*s/F) compared to Miss trials (0.42 ± 0.18 dF*s/F, p < 0.05), False Alarm trials (0.42 ± 0.12 dF*s/F, p < 0.001), and Correct Reject trials (0.66 ± 0.21 dF*s/F, p < 0.001) across animals and odor-pairs (Figure 2F). At the same time, in both Hit and False Alarm trials, HDB acetylcholine rapidly increased leading up to beam breaks at the reward port. Quantifying the slopes of the traces after odor delivery revealed that Hit trials exhibited steeper slopes (0.63 ± 0.12 dF/F/s) than Miss trials (0.23 ± 0.06 dF/F/s, p < 0.01), False Alarm trials (0.44 ± 0.07 dF/F/s, p < 0.05), and Correct Reject trials (0.22 ± 0.06 dF/F/s, p < 0.001). However, slopes in False Alarm trials were also significantly steeper compared to Miss (p < 0.05) and Correct Reject trials (p < 0.001) (Figure 2G). Together, these data revealed sharp increases in basal forebrain cholinergic tone which corresponded to reward-seeking behavior in both Hit and False Alarm trials. However only subsequent reward delivery in Hit trials led to a large, slow suppression of HDB cholinergic tone.

### 2.3 Reward-seeking and reinforcement-linked patterns of HDB cholinergic signaling are independent of learning

We next questioned whether patterns of cholinergic signaling in the HDB changed over the course of new odor-reward association learning. To determine this, we selected experiments with slower rates of learning, in which at least 3 blocks were performed with < 70% success, but criteria for learning were eventually met within 300 trials (N = 16 sessions, 5 animals). This paradigm allowed us to compare, within a testing session, cholinergic signaling from prelearning blocks (blocks with < 70% accuracy), to responses after an odor association had been effectively learned (first two consecutive blocks and subsequent blocks > 85% accuracy) (Figure 3A). Comparing pre-learning vs. learned blocks revealed similar patterns of cholinergic signaling in the HDB during both reward-seeking and after reward delivery in both Hit and False Alarm trials (Figure 3B). In agreement with the data from whole sessions (Figure 2), we found that the magnitude of the reward-related suppression in learned blocks was larger in Hit (1.60 ± 0.54 dF*s/F) than in False Alarm trials (0.57 ± 0.36 dF*s/F, p < 0.05). However, there was no difference in the magnitude of the reward-related suppression between pre-learning (Hit = 1.48 ± 0.42 dF*s/F, False Alarm = 0.47 ± 0.20 dF*s/F) compared to learned blocks in either Hit (p = 0.96) or False Alarm trials (p = 0.99) (Figure 3C). Additionally, across animals and odor pairs, the slope of the cholinergic signal after odor presentation was the same between pre-learning (Hit = 1.31 ± 0.32 dF/F/s, False Alarm = 0.90 ± 0.17 dF/F/s) and learned blocks (Hit = 1.41 ± 0.40 dF/F/s, False Alarm = 0.81 ± 0.19 dF/F/s) for both Hit (p = 0.84) and False Alarm trials (p = 0.89) (Figure 3D). These data suggest that HDB cholinergic signaling increases during reward-seeking behavior and decreases with reward delivery, regardless of whether the odor-reward association has been effectively learned. These data, however, do not address whether patterns of HDB cholinergic signaling drive the formation of an odor-reward association, or depend on the context of an odor reward association.

**Figure 3:**
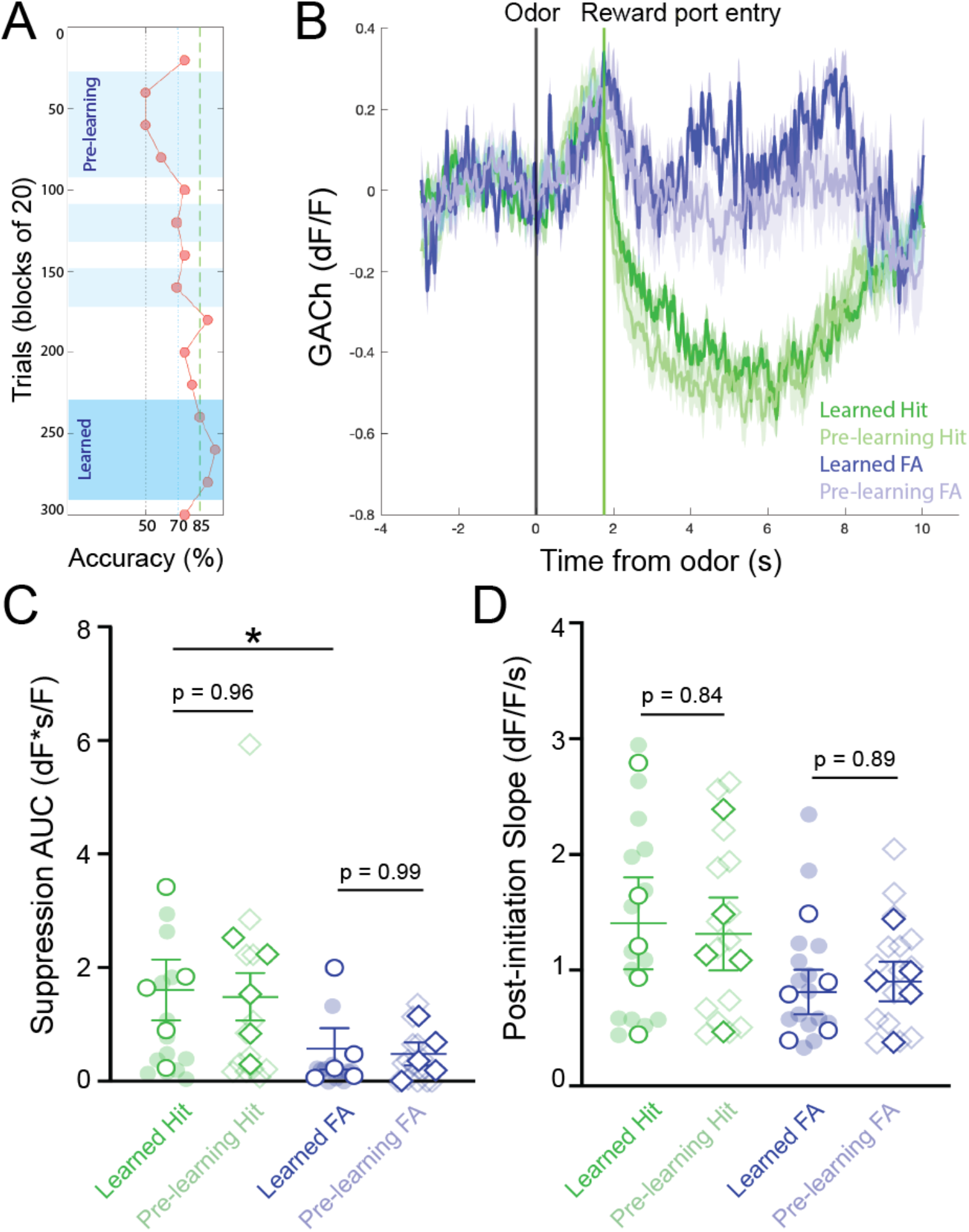
Temporal profile of cholinergic signaling in the HDB does not change with within-session discrimination learning. **A.** Accuracy in blocks of 20 trials for a go/no-go testing session highlighting blocks analyzed as “pre-learning” (light blue shading) and “learned” (dark blue shading). **B.** Average GACh dF/F traces for Hit and False Alarm trials separated into prelearning and learned trials. Shaded areas represent 95% confidence intervals. Black line marks trial initiation time. Green line marks the average reward port entry time. **C.** Area under the curve of suppression below baseline across trial types in pre-learning and learned blocks. Transparent circles and diamonds represent individual testing sessions. Hollow opaque circles and diamonds represent mean values from all sessions completed by individual mice. Lines and error bars show mean ± SEM of means from each animal. *p < 0.05, two-way nested repeated measures ANOVA with Tukey correction for multiple comparisons. **D.** Slopes of GACh dF/F after trial initiation across trial types in pre-learning and learned blocks. Transparent circles and diamonds represent individual testing sessions. Hollow opaque circles and diamonds represent mean values from all sessions completed by individual mice. Lines and error bars show mean ± SEM of means from each animal. Lines and error bars show mean ± SEM of means from each animal. Two-way nested repeated measures ANOVA with Tukey correction for multiple comparisons.

### 2.4 Reward related suppression of HDB cholinergic signaling is task-dependent and relies on association between odor cue and reward

We next sought to determine whether patterns of basal forebrain cholinergic signaling were influenced by the context of the olfactory task, including the requirement for odor discrimination and the reliable association between odor and reward. An alternative possibility was that the observed pattern in HDB cholinergic signaling may simply reflect reward-seeking and consumption behaviors, independent of odor discrimination or odor-reward association. While task dependence would suggest that HDB cholinergic signaling is involved in top-down regulation, the latter possibility would suggest that HDB cholinergic signaling responds to bottom-up cues. To distinguish between these possibilities, we recorded basal forebrain cholinergic signals during a version of the go/no-go task, in which there was no association between the odor presented and the availability of the reward (pseudo-learning, N = 10 sessions, 4 animals). In the pseudo-learning paradigm, S+ and S− odors were each presented 50% of the time, and a water reward was available on 50% of the trials at random (Figure 4A). This version of the task retains odor presentations, odor detection, reward-seeking, and reward delivery, removing only the odor-reward association and the need for odor discrimination. During pseudo-learning, mice typically obtained ~50% success rate with a mix of trial types biased toward positive, reward-seeking responses (Pseudo FA and Pseudo Hit trials), and against negative (Reject) responses (Figure 4B).

**Figure 4:**
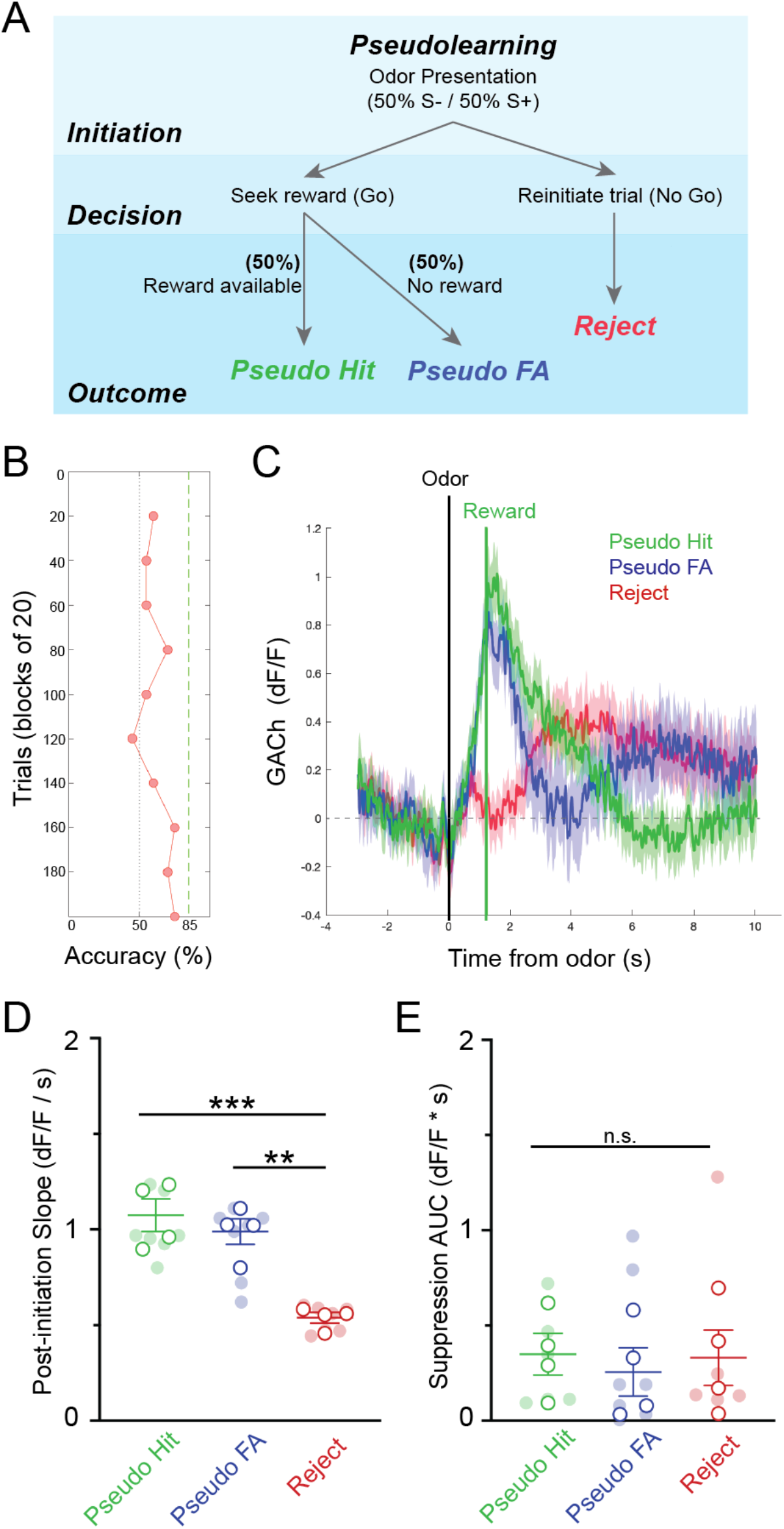
Pseudo-learning reveals task-dependence of dynamic basal forebrain cholinergic tone. **A.** Schematic of olfactory-cued go/no-go pseudo-learning task showing odor presentation, decisions, and possible trial outcomes (Pseudo-Hit, Pseudo-False Alarm, Reject). **B.** Accuracy in blocks of 20 trials for a pseudo-learning testing session showing performance near chance. **C.** Average GACh dF/F traces for Pseudo-Hit, Pseudo-False Alarm and Reject trials. Shaded areas represent 95% confidence intervals. Black line marks trial initiation time. Green line marks the average reward port entry time in reward-seeking trials. **D.** Average GACh dF/F traces for each trial type. Shaded areas represent 95% confidence intervals. Black line marks trial initiation time. Green line marks the average reward port entry time in Hit and False Alarm trials. **E.** Slopes of GACh dF/F after trial initiation across trial types and testing sessions. Transparent circles represent individual testing sessions. Hollow circles represent mean values from all sessions completed by individual mice. Lines and error bars show mean ± SEM of means from each animal. **p < 0.01, *** p < 0.001, two-way nested repeated measures ANOVA with Tukey correction for multiple comparisons. **F.** Area under the curve of suppression below baseline across trial types and testing sessions. Transparent circles represent individual testing sessions. Hollow circles represent mean values from all sessions completed by individual mice. Lines and error bars show mean ± SEM of means from each animal. Two-way nested repeated measures ANOVA with Tukey correction for multiple comparisons.

Averaging across trials of the pseudo-learning task showed similar patterns of basal forebrain cholinergic signaling between Pseudo-Hit and Pseudo-FA trials, both of which differed from Reject trials (Fig 4C). In both Pseudo-Hits and Pseudo-FA trials, cholinergic signaling increased after odor presentation as mice seek rewards. This was reflected in significantly shallower slopes after trial initiation for Reject trials (0.54 ± 0.03 dF/F/s), compared to PseudoHit trials (1.07 ± 0.09 dF/F/s, p < 0.001) and Pseudo-FA trials (0.99 ± 0.07 dF/F/s, p < 0.01) (Figure 4D), suggesting that increased HDB acetylcholine reflects reward-seeking behavior independent of an odor-reward association. Strikingly, however, we did not observe a slow suppression in cholinergic tone following reward delivery in Pseudo-Hit trials. The total magnitude of suppression was not larger in Pseudo-Hit trials (0.35 ± 0.12 dF*s/F) compared to Pseudo-FA (0.26 ± 0.13 dF*s/F, p = 0.96) or Reject trials (0.33 ± 0.15 dF*s/F, p = 0.98) (Fig 4E). Together, these data indicate that the reward-related suppression of basal forebrain cholinergic tone does not merely reflect reward delivery or reward consumption. Rather, the suppression depends on the task context and requires an association between odor and reward.

### 2.5 HDB GABAergic neuronal activity mirrors cholinergic tone in response to positive reinforcement

Having revealed dynamic cholinergic signaling in the HDB during olfactory-guided behavior, we next examined a potential target of local cholinergic signaling. Basal forebrain GABAergic neurons express both metabotropic and nicotinic acetylcholine receptors, and have been shown to respond to local cholinergic signaling (Yang et al., 2014; Xu et al., 2015). We hypothesized that HDB GABAergic neuronal activity would be controlled, in part, by local cholinergic signaling and that GABAergic neuronal activity would follow a similar pattern of activation and suppression across phases of go/no-go task performance. To test this, we selectively expressed GCaMP in HDB GABAergic neurons by injecting an AAV encoding cre-dependent GCaMP6M (AAV flex-GCaMP6M) into Vgat-Cre mice (Figure 5A, B). We then recorded GABAergic neuronal activity via fiber photometry during performance of the go/no-go task. After behavioral shaping, new odor learning was accomplished within sessions of 200-300 trials (Figure 5C). Aligning individual trials by trial initiation time revealed bidirectional modulation of GABAergic neuronal activity with excitation following odor deliver and suppression following reward delivery in Hit trials (Figure 5D). Similar to the changes we observed in cholinergic tone, HDB GABAergic neuronal activity was reliably suppressed following reward delivery. Areas under the curve of the suppression below baseline were significantly larger in Hit trials (8.36 ± 1.25 dF*s/F) than in False Alarm (1.69 ± 0.48 dF*s/F, p < 0.001), Correct Reject (0.85 ± 0.30 dF*s/F, p < 0.001), or Miss trials (1.72 ± 0.94 dF*s/F, p < 0.001) (Figure 5E). In contrast to changes in cholinergic tone, however, HDB GABAergic neurons responded to both the S+ and S− odors. This was reflected in positive slopes of the GCaMP signal following trial initiation in Hit (1.67 ± 0.33 dF/F/s, p < 0.05), Miss (1.61 ± 0.26 dF/F/s, p < 0.01), False Alarm (1.64 ± 0.32 dF/F/s, p < 0.05), and Correct Reject trials (1.33 ± 0.24 dF/F/s, p = < 0.05) (Figure 5F). Additionally, slopes from different trial types were not significantly different from each other (p = 0.34). The consistent response to both odors implied that basal forebrain GABAergic neurons receive bottom-up olfactory information, and that their activity may reflect odor detection or active sensing.

**Figure 5:**
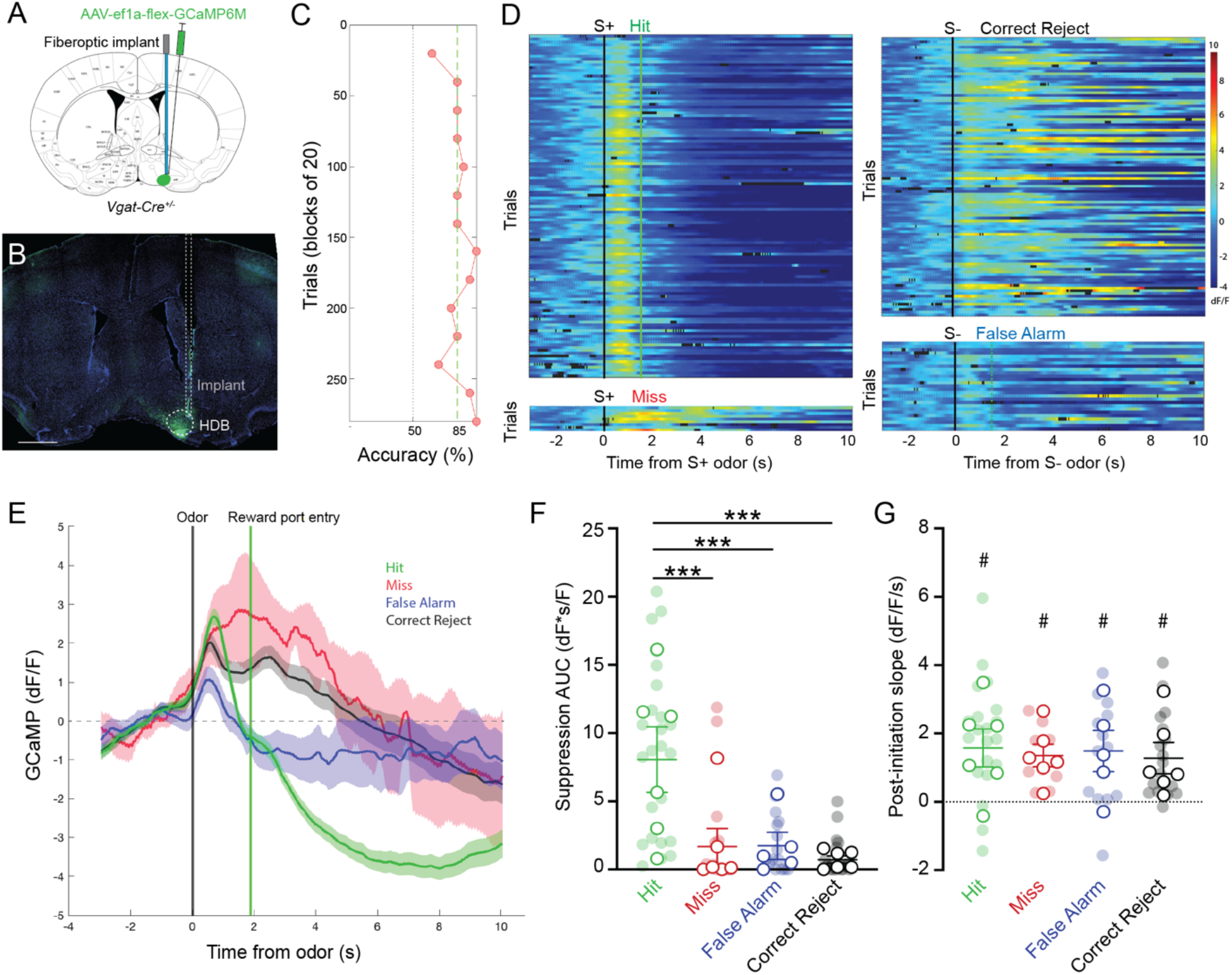
HDB GABAergic neuronal activity mirrors cholinergic tone in response to positive reinforcement **A**. Coronal section schematic showing AAV injection and implant targeting the HDB. **B.** IHC of a coronal section showing GCaMP6M expression and implant targeting in the HDB. Scale bar = 1 mm. **C.** Accuracy in blocks of 20 trials for a go/no-go testing session with novel odors highlighting chance (50%) and criteria (85%) levels. Accuracy is from the same session as the trials shown in D and E. D. **D.** Heatmap showing isosbestic-subtracted GCaMP dF/F from individual trials in a single go/no-go testing session. Trials are aligned by trial initiation time and divided by trial outcome. **E.** Average GCaMP dF/F traces for each trial type in the session shown in D. Shaded areas represent 95% confidence intervals. Black line marks trial initiation time. Green line marks the average reward port entry time in Hit and False Alarm trials. **F.** Area under the curve of suppression below baseline across trial types and testing sessions. Transparent circles represent individual testing sessions. Hollow circles represent mean values from all sessions completed by individual mice. Lines and error bars show mean ± SEM of means from each animal. *** p < 0.001 two-way nested repeated measures ANOVA with Tukey correction for multiple comparisons. **G.** Slopes of GCaMP dF/F after trial initiation across trial types and testing sessions. Transparent circles represent individual testing sessions. Hollow circles represent mean values from all sessions completed by individual mice. #p < 0.05, ##p < 0.01, nested one sample t test comparing trial type values to 0. Two-way nested repeated measures ANOVA with Tukey correction for multiple comparisons shows p = 0.36 for differences between trial types.

## 3 Discussion

Sensory perception relies on a combination of bottom-up sensory input, and top-down behavioral state-dependent regulation. The basal forebrain serves as a key mediator of top-down regulation related to the behavioral states of attention, arousal, and wakefulness (Muir et al., 1993; Voytko et al., 1994; Szymusiak, 1995; Sarter and Bruno, 1999; Hasselmo and McGaughy, 2004; Herrero et al., 2008; Goard and Dan, 2009; Anaclet et al., 2015; Kim et al., 2015; Zant et al., 2016). Many of these effects are thought to be mediated by cholinergic signaling at downstream sensory circuits. Supporting this, in olfactory, visual, and auditory circuits, cholinergic neuromodulation has been shown to increase gain, improve signal to noise ratios, increase pattern separation, and increase the weight of bottom-up sensory input (Mandairon et al., 2006; Herrero et al., 2008; Chaudhury et al., 2009; Ghatpande and Gelperin, 2009; Goard and Dan, 2009; Ma and Luo, 2012; Chapuis and Wilson, 2013; Zhan et al., 2013; Rothermel et al., 2014). However, mounting evidence suggests that parallel GABAergic outputs from the basal forebrain also play a significant role sculpting downstream circuit activity (Nunez-Parra et al., 2013, 2020; Kim et al., 2015; Xu et al., 2015; Böhm et al., 2020; Villar et al., 2020).

In olfaction, input from the basal forebrain significantly impacts the earliest stages of signal transduction in the olfactory bulb. Separately, the cholinergic and GABAergic projection pathways from the HDB drive distinct changes in olfactory bulb neuronal activity. For example, GABAergic projections from the HDB synapse onto inhibitory granule cells and periglomerular interneurons in the olfactory bulb where they mediate disinhibition and desynchronization of Mitral Cell firing bulb (Gracia-Llanes et al., 2010; Sanz Diez et al., 2019; Villar et al., 2020). Moreover, experiments implementing chemogenetic inhibition showed that basal forebrain GABAergic projections are required for effective odor discrimination (Nunez-Parra et al., 2020). On the other hand, other experiments have revealed that basal forebrain cholinergic projections to the olfactory bulb increase excitability, modulate signal to noise ratios in mitral cell firing, and can rapidly dishabituate olfactory bulb odor responses (Ma and Luo, 2012; Rothermel et al., 2014; Ogg et al., 2018). Finally, more recent studies have directly compared optogenetic stimulation of basal forebrain cholinergic and GABAergic terminals within the olfactory bulb (Böhm et al., 2020), describing that local stimulation of cholinergic terminals increased mitral cell firing during sniffing regardless of odor presentation, whereas stimulation of GABAergic terminals decreased spontaneous mitral cell firing and increased firing during sniffing only when odors were presented. Together, these findings imply that basal forebrain cholinergic and GABAergic neurons mediate distinct features of top-down regulation, and that both types of basal forebrain projections modulate olfactory bulb odor and sniff responses.

Importantly, cholinergic and GABAergic neurons in the basal forebrain work together to modulate downstream circuit function (Dannenberg et al., 2015; Böhm et al., 2020). An outstanding question is how parallel cholinergic and GABAergic output pathways are coordinated during behavior at the level of the basal forebrain. In the current study, we show that cholinergic signaling within the basal forebrain is dynamically regulated during olfactory-guided behavior. We also show that distinct changes in basal forebrain cholinergic tone correspond to changes in neighboring GABAergic neuronal activity. These results suggest that cholinergic signaling within the basal forebrain may dynamically drive behavior and behavioral statedependent changes in the basal forebrain output pathways that mediate top-down regulation of olfactory processing.

### 3.1 Bidirectional changes in basal forebrain cholinergic tone during olfactory task performance

Though basal forebrain neuronal activity has been characterized across a variety of behavioral states and in a number of sensory discrimination and association learning tasks (Mandairon and Linster, 2009; Devore et al., 2015; Hangya et al., 2015; Xu et al., 2015; Harrison et al., 2016; Nunez-Parra et al., 2020), how this activity is regulated by local signaling within the basal forebrain remains largely unknown. To investigate local cholinergic signaling within the basal forebrain we used a GPCR Activation-Based sensor for acetylcholine (Jing et al., 2018) combined with fiber photometry. This approach allowed us to record changes in acetylcholine from the basal forebrain with sub-second temporal resolution, in freely behaving animals.

With this approach we were able to record rapid changes in cholinergic tone from the HDB during free exploration of an open field arena, and during olfactory-cued operant behavior. Basal forebrain neuronal activity has been previously correlated with locomotion and slow changes in arousal (Sarter and Bruno, 1999; Goard and Dan, 2009; Xu et al., 2015). However, while we observed frequent spontaneous activation and suppression signaling events during exploration, significant changes in GACh fluorescence were not correlated with locomotion. This highlights an interesting discrepancy between cholinergic neuron activity and the local cholinergic signaling itself. Our data indicate that basal forebrain acetylcholine changes on a rapid timescale, which was not solely reflective of slow changes in behavioral state. Instead, changes in basal forebrain cholinergic tone were temporally precise based on behavioral action. This raises the possibility that changes in HDB cholinergic tone dynamically control basal forebrain output on a moment-to-moment basis, in line with the performance of complex olfactory guided behaviors.

To examine cholinergic signaling dynamics during complex olfactory guided behavior, we tested mice on a freely moving, olfactory-cued go/no-go discrimination task. The task included self-initiation of trials, followed by periods of active sensing, odor detection, discrimination, reward-seeking, and positive / negative reinforcement. We hypothesized that cholinergic signaling would be dynamically regulated within trials of the go/no-go task, reflecting changing needs for basal forebrain mediated top-down regulation during different behaviors, and in response to reinforcement. Supporting this hypothesis, we observed rapid, bidirectional changes in acetylcholine that were time-locked to phases of the go/no-go task. Specifically, we found that acetylcholine increased rapidly in the basal forebrain during reward-seeking behavior. Once the availability of a reward was ascertained, acetylcholine responses decreased rapidly. Finally, if a reward was successfully obtained, acetylcholine decreased slowly, but transiently, below baseline levels. Recent studies reported decreased activity in a subset of basal forebrain neurons following both stimulus presentation and reward delivery (Nunez-Parra et al., 2020). However, in a small population of identified cholinergic neurons, no reliable changes in firing rate were detected with reward delivery. This discrepancy may further suggest a disconnect between neuronal activity and local cholinergic tone. Alternatively, these data may reflect differences in the task requirements between our freely moving task and previously described head-fixed experiments (Nunez-Parra et al., 2020). Ultimately, the complex cholinergic signaling dynamics that we observed suggest that local cholinergic signaling corresponds to distinct features of the go/no-go task, perhaps reflecting reward-seeking behavior and subsequent reward delivery. The reward-related suppression of cholinergic signaling in Hit trials was particularly interesting given the implication that a baseline cholinergic tone in the basal forebrain is selectively suppressed in response to positive feedback.

### 3.2 Task-dependent and learning-independent patterns of cholinergic signaling in the basal forebrain

These observations led us to question whether cholinergic signaling dynamics in the basal forebrain were (1) a driver of odor-reward association learning, (2) a consequence of association learning, or (3) independent of odor-reward association, and instead linked to the performance of task-related behaviors. To directly investigate these possibilities, we examined cholinergic signaling dynamics over the course of odor-reward association learning, and in the absence of odor-reward associations. If task-linked cholinergic signaling dynamics are a consequence of odor-reward association learning, we might have expected temporal profiles of cholinergic signaling to change over the course of single sessions, where new odor-reward associations are being learned. For example, reward expectation scales with increasing success over a go/no-go session as new odor-reward associations are effectively learned (Tremblay et al., 1998). Thus, if increased cholinergic signaling in the basal forebrain reflects reward expectation, we would expect reporter responses to change over the course of learning within sessions. Indeed, a recent study recording neuronal activity in the basal forebrain found that a higher percentage of cholinergic and non-cholinergic neurons changed their firing rates in response to an odor cue after an odor-reward association was learned in a go/no-go task (Nunez-Parra et al., 2020). However, examining cholinergic tone directly, we find that changes in acetylcholine during go/no-go trials are stable over the course of odor-reward association learning. Neither increased acetylcholine during reward-seeking, nor suppressed acetylcholine following reward delivery, change over the course of a learning session. Intriguingly, rapid changes in basal forebrain cholinergic tone during reward-seeking did not reflect the strength of reward expectation. Such stability of basal forebrain cholinergic signaling over the course go/no-go testing sessions suggests that acetylcholine release within the basal forebrain is either an upstream driver of learning - relating specific perceptual decisions to positive and negative outcomes - or it may be all together independent of odor-reward associations - reflecting only reward-seeking and reward-consumption behaviors.

To distinguish between these possibilities, we examined cholinergic signaling during pseudo-learning, a version of the go/no-go task in which rewards were randomly available 50% of the time, regardless of the odor presented. Pseudo-learning preserves odor detection (the animals can only seek a reward after receiving an odor presentation), reward-seeking, and reward delivery, but removes the association between odor and reward. If basal forebrain cholinergic signaling is simply a reflection of reward-seeking and consumption, we would have expected to observe the same stable patterns of cholinergic signaling with reward-seeking and reward delivery that we observed in the go/no-go discrimination task. Alternatively, if basal forebrain cholinergic signaling serves a role in relating the odor-discrimination context of the task to reward-seeking behavioral choices or positive reinforcement outcomes, we would expect to observe differences when the rules of the task are changed. Indeed, in the pseudo-learning task, removing the cue-reward association led to a decrease in the reward-related suppression of basal forebrain cholinergic tone. However, increased cholinergic tone during reward-seeking behavior occurred regardless of odor-reward association. Surprisingly, patterns of basal forebrain cholinergic signaling were stable over the course of pseudo-learning sessions. It’s possible that this was because mice quickly realized that the context of the task had changed and rapidly altered their strategy to fit the new rules of the pseudo-learning task. If so, the reward-related suppression of cholinergic signaling may not only reflect reward delivery but also take into account knowledge of the task itself.

### 3.3 Odor-evoked activity and reward-related suppression of basal forebrain GABAergic neurons during olfactory task performance

GABAergic neurons in the HDB express cholinergic receptors and respond to local acetylcholine release (Yang et al., 2014; Xu et al., 2015). At the same time, GABAergic output from the HDB mediates distinct forms of top-down regulation important for state-dependent active sensing and odor discrimination (Nunez-Parra et al., 2013; Böhm et al., 2020). Having found that local cholinergic tone is dynamically regulated during performance of an olfactory-cued go/no-go task, we next examined whether changes in local cholinergic tone corresponded to changes in basal forebrain GABAergic neuronal activity. We reasoned that if HDB GABAergic neuronal activity is dynamically controlled by local cholinergic signaling, we might expect to observe correlations between temporal profiles of GABAergic neuronal activity and cholinergic tone during olfactory discrimination tasks. However, in contrast to the observed cholinergic signaling patterns, we observed GABAergic responses to both the S+ and S− odors across all trial types, regardless of reward-seeking behavior. Intriguingly, recent studies report that stimulating GABAergic projections to the olfactory bulb enhanced sniff-locked odor responses from a subset of mitral cells, while suppressing spontaneous activity (Böhm et al., 2020). Inhibiting these projections, on the other hand, reduced odor discrimination (Nunez-Parra et al., 2013). In this context, our data show that HDB GABAergic neurons respond broadly during odor discrimination, and they suggest that they may mediate enhanced odor discrimination during active sniffing.

At the same time, we observed similar changes in cholinergic tone and GABAergic neuronal activity in response to positive reinforcement. Both GABAergic neuronal activity and cholinergic tone were suppressed following reward delivery in Hit trials. Suppression below baseline implies that a population of GABAergic neurons in the basal forebrain are tonically active. Notably, tonic and rhythmic neuronal firing has been observed in basal forebrain, and is strongly dependent on behavioral state (Nunez, 1996; Détári et al., 1999; Szymusiak et al., 2000). The similarity between the suppression of basal forebrain GABAergic neuronal activity and local cholinergic signaling following reward delivery suggests that activity of basal forebrain GABAergic neurons is influenced by local cholinergic tone. Importantly however, our data do not distinguish whether basal forebrain GABAergic neurons are a target of tonic excitement from local acetylcholine. Indeed, other studies have reported that cholinergic collateralization within the basal forebrain directly activates local non-cholinergic and/or GABAergic neurons (Yang et al., 2014; Dannenberg et al., 2015; Xu et al., 2015; Nunez-Parra et al., 2020). In this context, our data raise the possibility that higher ambient cholinergic tone and increased tonic activity of basal forebrain GABAergic neurons in awake states create an environment where such signals can be bidirectionally modulated to solidify learned cue-reward associations.

Here we have revealed rapid, bidirectional changes in cholinergic tone within the basal forebrain during complex, olfactory-guided behavior. Characterization of the cholinergic signal itself through visualization of the GACh reporter sigal is a first step towards understanding how cholinergic drive influences basal forebrain circuitry, and thus, top-down regulation of sensory processing. Future work will be needed to determine the mechanistic impact of dynamic cholinergic tone on specific HDB projection neuron populations. The current data, however, support the idea that local HDB cholinergic signaling is dynamically regulated by behavior and behavioral state, making it an intriguing candidate for coordinating state-dependent effects on HDB circuits and projection outputs.

## 4 Materials and Methods

### 4.1 Animals

Mice were maintained on a 12 h light-dark cycle and were treated in compliance with the US Department of Health and Human Services and Baylor College of Medicine IACUC guidelines. C56Bl6/J and Vgat-cre male and female mice underwent surgery at 2–4 months old. Vgat-Cre (*Slc32a1*tm2(cre)Lowl, Stock: 028862) mice were originally purchased from Jackson Laboratories.

### 4.2 Surgical Procedures

Mice were anesthetized with 4% isoflurane in O2 and maintained under anesthesia with 1–2% isoflurane in O2. Craniotomies were made over the sites of stereotaxic injections and fiberoptic implants that were guided by Angle Two software (Leica) normalized to Bregma. To target viral expression to the HDB, a unilateral injection of virus was made into the left HDB (from Bregma: ML −1.0 mm, AP 0.1 mm, DV −5.45 mm). Viruses were packaged in-house and included AAV-hsyn-GACh2.0, Serotype DJ8 injected into C57Bl6/J (WT) mice and AAV-ef1α-flex-GCaMP6M, Serotype DJ8, injected into Vgat-cre mice. The plasmid containing GACh2.0 was a generous gift from the Yulong Li Lab (Jing et al., 2018). 250 nL of virus was injected into the HDB over ten minutes. Following viral injection, the injection needle was removed and a custom fiber optic implant (0.48 na, 200 um core diameter, RWD systems) was lowered to a target 0.1 mm dorsal to the injection target. The implant was then fixed in place with Metabond dental cement (Parkell). Mice were allowed to recover for 3 weeks before behavioral experiments. HDB targeting was verified in all cases with immunofluorescence imaging of the implant track and viral expression within the HDB.

### 4.3 Go/no-go behavior

Prior to photometric recording, mice underwent behavioral shaping, allowing them to learn the mechanics of the go/no-go task. Mice progressed through 5 behavioral shaping stages over the course of 10-14 days as described previously (Quast et al., 2016; Liu et al., 2017). Briefly, mice were water-restricted to no less than 85% of their baseline weight for 2 d before shaping. Water was restricted to 40 mL per kg, per day during the restriction period. Mice trained using a go/no-go paradigm in a behavioral chamber with infrared nose pokes (Med Associates Inc.). All mice were first trained to poke their nose into the odor port for at least 300 ms, before moving to the side water port to retrieve water reward within 5 s (Figure 2). After preliminary training sessions (~30–60 min/d for ~5–6 d) mice were trained to respond to the S+ odor cue (1% Eugenol in mineral oil) by moving to the water port for a reward and were trained to respond to the S− odor (1% Methylsalicylate in mineral oil) by refraining from poking into the reward port and, instead, initiating a new trial. We required mice to sample odors for at least 100 ms before responding and to respond within 5 s after trial initiation. False alarms (incorrect response to S− odor) caused a 4 s timeout punishment. S+ and S-stimuli (Table 1) were presented to the mice in random sequences during training. Mice were trained for 20 trials per block and ~10-15 blocks per day. Throughout shaping and testing accuracy was calculated by block of 20 trials. Odors pairs were considered “learned” after two consecutive blocks > 85% accuracy. After 3–6 d of odor training, the mice performed at over 85% correct responses. Mice were then tested on new odors (Table 1) diluted to 1% in mineral oil during photometric recording.

**Table 1:** Monomolecular odors used in go/no-go shaping and testing

| *Odorant* | *Molar Mass*<br>*(g/mol)* | *Vapor*<br>*Pressure*<br>*(mmHg)* | *Functional Group* |
| --- | --- | --- | --- |
| *(-) Carvone* | 150.22 | 0.16 | Cyclic ketone, 10C, 1 double bond |
| *(+) Carvone* | 150.22 | 0.16 | Cyclic ketone, 10C, 1 double bond |
| *1-Butanol* | 74.12 | 7 | Straight chain alcohol, 4C |
| *1-heptanol* | 116.2 | 0.2163 | Straight chain alcohol, 7C |
| *1-Hexanol* | 102.18 | 1 | Straight chain alcohol, 6C |
| *1-Pentanol* | 88.15 | 44.6 | Straight chain alcohol, 5C |
| *Acetophenone* | 120.15 | 0.397 | Aromatic ketone |
| *alpha-Pinene* | 136.24 | 3 | Cyclic, 7C, 1 double bond |
| *Citral* | 152.24 | 0.22 | Aldehyde, 10C |
| *Ethyl Acetate* | 88.11 | 93.2 | Ester, 4C |
| *Eucalyptol* | 154.249 | 1.9 | Cyclic ether, monoterpinoid |
| *Eugenol* | 164.2 | 0.0221 | Aromatic alcohol, ether |
| *Isoamyl acetate* | 130.19 | 4 | Ester, 7C |
| *(-) Limonene* | 150.22 | 0.16 | Cyclic monoterpene, 10C, 2 double bonds |
| *(+) Limonene* | 150.22 | 0.16 | Cyclic monoterpene, 10C, 2 double bonds |
| *Menthone* | 154 | 0.895 | Cyclic ketone, 10C, no double bonds |
| *Methyl Acetate* | 74.08 | 173 | Ester, 3C |
| *Methyl Salicylate* | 152.15 | 0.0343 | Aromatic alcohol, ester |

For pseudo learning (Figure 4), S+ and S− odors were presented randomly, each 50% of the time, as in the go/no-go discrimination task. Reward availability was also randomly determined with rewards available upon reward-port entry 50% of the time.

### 4.4 Photometry

To allow stimulation and recording of fluorescent transients through the same fiberoptic implant, we utilized a fiber photometry system from Doric lenses. Two light emitting diodes (465 nm and 405 nm wavelength) were coupled to a filter cube by 0.48 na, 400 um core diameter fiber optic cables. The filter cube separated excitation and emission wavelengths, directing the excitation wavelengths along a 0.48 na, 200 um core diameter fiber optic toward the mouse through a rotary connector attached to the behavior box. Emission wavelengths were carried from the mouse to the filter cube along the same fiber, then directed to a femtowatt photodetector (Newport) through a 0.48 na, 600 um core diameter fiber optic cable. Excitation and emission were controlled and recorded respectively in Doric Studio software. Both GACh2.0 and GCaMP6M were excited at 465 to record either acetylcholine (for GACh2.0) or calcium binding (for GCaMP). Additionally, excitation at the isosbestic point for GCaMP (405 nm) generates emission which is insensitive to calcium binding. Thus, a photometric recording of GCaMP with excitation at 405 nm is a useful control for motion artifacts and other calcium-independent noise. For GACh2.0, the isosbestic point is near 405 allowing it to serve as a control signal in a similar manner. To record from the control channel (excited at 405 nm) and the experimental channel (excited at 465 nm) simultaneously, we employed a “locked-in” strategy where each LED was modulated at a different high frequency. Emission resulting from both modes of excitation was recorded by the same photodetector and the signal was demodulated online in Doric Studio to separate the control channel form the experimental channel. Both signals were then converted to df/f and the control channel was subtracted from the experimental channel to reduce noise.

### 4.5 Histology

For immunohistochemistry, mice were deeply anesthetized then transcardially perfused with PBS followed by 4% PFA. Brains were removed and immersion fixed in 4% PFA overnight at 4°C. Brains were transferred to 30% sucrose and allowed to equilibrate, then they were frozen and sectioned at 40 μm on a cryostat (Leica). The sections were washed in 0.3% PBS-T, then incubated in a blocking solution composed of 10% normal goat serum, 0.3% PBS-T, and 3M glycine for 1 h at room temperature or overnight at 4°C. Following blocking, slices were incubated in chicken ∝ GFP primary antibody (1:1,000, Abcam, ab13970) diluted in blocking buffer overnight at 4°C. The next day slices were washed 3× in 0.3% PBS-T then incubated in Goat ∝ Chicken:488 secondary antibody (1:1,000, Invitrogen, A32931) for 2 h at room temperature. Slices were then washed 3× in 0.3% PBS-T with Hoescht included in the middle wash. After the final wash slices were transferred to 0.5× PBS and mounted on glass slides with glycerol-based mounting media (Southern Biotech). Slices were imaged on a Leica SP8 Confocal with 10× air objectives.

### 4.6 Statistics and Data Analysis

Isosbestic-subtracted dF/F traces were extracted and segmented according to the timing of IR beam breaks during go/no-go behavior using custom MATLAB scripts. Photometry traces for individual trials were separated by trial outcome and averaged within sessions. Importantly, we do not apply trial-to-trial or trial-averaged baseline subtraction or amplitude normalization. Thus, values reported reflect isosbestic-subtracted dF/F. 95% confidence intervals were calculated within sessions using traces from individual trials. For learning-related analyses (Figure 3) sessions with 3 or more blocks < 70% accuracy and with 2 or more consecutive blocks > 85% accuracy were sub-selected from the larger dataset. Hit and False Alarm trials from blocks < 70% accuracy were grouped and analyzed separately as “pre-learning” trials. Hit and False Alarm trials from the first two consecutive blocks > 85% accuracy and from subsequent blocks > 85% accuracy were grouped and analyzed separately as “learned” trials. In all cases, postinitiation slopes were calculated by linearly fitting the data after trial initiation before. Areas under the curve were calculated by summing negative values of average traces after trial initiation and dividing by sampling rate. In all cases, comparisons of post initiation slopes and areas under the curve between trial types utilized a nested, two-way, repeated measures ANOVA. This analysis maintains the relationship between trial types within a single session (repeated measures). Additionally, nesting multiple sessions recorded from the same animal provides a conservative statistical measure which consider all sessions from a single animal together. In the case of post-initiation slopes of GCaMP traces from GABAergic neurons, values were also compared to 0 using a nested one-sample t test. All reported values reflect means from the nested analyses ± SEM and in all cases p < 0.05 was considered significant.

## 6 Author Contributions

EH designed and conducted experiments, analysed data, and wrote the manuscript. KBA conducted experiments, analysed data, and helped write and edit the manuscript. BRA acquired funding, provided guidance on experimental design, data analysis, and interpretation, and helped edit the manuscript.

## 8 Acknowledgments

We would like to thank the Postbaccalaureate Research Education Program (PREP) at Baylor college of medicine for their support of KBA, the neurobehavioral core at Baylor College of Medicine for sharing behavioral equipment, and the Arenkiel lab for many helpful discussions.

